# Intermittent cell division dynamics in regenerating Arabidopsis roots reveals complex long-range interactions

**DOI:** 10.1101/2023.11.28.569038

**Authors:** T. Fallesen, S. Amarteifio, G. Pruessner, H. J. Jensen, G. Sena

## Abstract

In this work, we present a quantitative comparison of the cell division dynamics between populations of intact and regenerating root tips in the plant model system *Arabidopsis thaliana*. To achieve the required temporal resolution and to sustain it for the duration of the regeneration process, we adopted a live imaging system based on light-sheet fluorescence microscopy, previously developed in the laboratory. We offer a straightforward quantitative analysis of the temporal and spatial patterns of cell division events showing a statistically significant difference in the frequency of mitotic events and spatial separation of mitotic event clusters between intact and regenerating roots.

## INTRODUCTION

Tissue regeneration, or the re-establishment of the form and function of a damaged or lost structure, is an example of post-embryonic morphogenesis. The history of regeneration research is long and rich in breakthroughs (Dinsmore, 1991), and some of the key molecular and mechanical details have been understood in recent decades (Elchaninov et al., 2021; Ikeuchi et al., 2016; Liu et al., 2023; Morinaka et al., 2023; Sugimoto et al., 2019).

The role of cell proliferation in the re-establishment of lost structures has long been recognized as central to the process of regeneration (Morgan, 1901). At the most fundamental level, there are basic yet unanswered questions regarding the type of dynamics and the parameters controlling it. For example, does cell proliferation during regeneration follow unique dynamics, distinguished from the ones driving other types of morphodynamics such as embryonic development or post-embryonic organogenesis such as metamorphosis in animals or flower formation in plants? Is regeneration a smooth process or does it go through sharp transitions, perhaps analogous to phase transitions observed in many complex dynamical systems? Unfortunately, more than one hundred years after the first observations, a complete quantitative description of cell proliferation dynamics during organ regeneration is lacking, impeding our efforts to understand how biological shapes and functions are established and maintained.

Here, we present a quantitative analysis of cell divisions in regenerating root tips of the plant model system *Arabidopsis thaliana*. Given the relatively long duration of root regeneration following full tip excision (Sena et al., 2009), we adopted light-sheet microscopy for sustained, high-resolution, time-lapse imaging. In plants, this method had been previously adapted first to *Arabidopsis* roots (Maizel et al., 2011; Sena et al., 2011) and then to other tissues (Berthet & Maizel, 2016; Clark et al., 2020).

Quantitative analyses of cell divisions in intact, i.e., *Arabidopsis* roots have a long history of not regenerating. Modern imaging methods span from simple light microscopy (Beemster & Baskin, 1998) to confocal microscopy (Campilho et al., 2006; Lavrekha et al., 2017; Rahni & Birnbaum, 2019) and light-sheet microscopy (Balaguer et al., 2016; Buckner et al., 2019; Sena et al., 2011; Wangenheim et al., 2016), but no comparison has been attempted between these dynamics and those in regenerating roots.

Algorithms to track cell divisions in light-sheet microscopy 4D datasets have been developed multiple times (Amarteifio et al., 2021; Buckner et al., 2019; Sena et al., 2011). For this work, we adopted hardware and software previously developed in our lab (Amarteifio et al., 2021; Baesso et al., 2018).

By comparing the dynamics of cell proliferation in a growing intact root with that in a regenerating one, in this work, we address the following fundamental questions: Is there a quantitative difference between the dynamics of cell division in an uncut root and that in a regenerating one? Is there a clear transition between different “phases” in cell division dynamics during root regeneration?

## RESULTS

### The temporal sequence of mitotic events is intermittent

The cyclin-dependent protein kinase CYCB1;1 is commonly used as a reporter of the G2/M transition in the cell cycle and, indirectly, of mitotic events (Reddy et al., 2004). Transgenic *Arabidopsis* plants expressing CYCB1;1::GFP (Reddy et al., 2004) were mounted on an open hardware light-sheet microscope setup (Baesso et al., 2018) specifically designed for imaging and tracking a single root tip every 15 minutes (see Methods).

The raw images were processed using our previously published routine (Amarteifio et al., 2021) to track and count the mitotic events in 3D. The number of cell divisions detected in each frame follows an intermittent temporal pattern with a noisy baseline below 20 events per frame punctuated by a few isolated bursts of much higher activity (Fig. 1).

**Figure 1.**
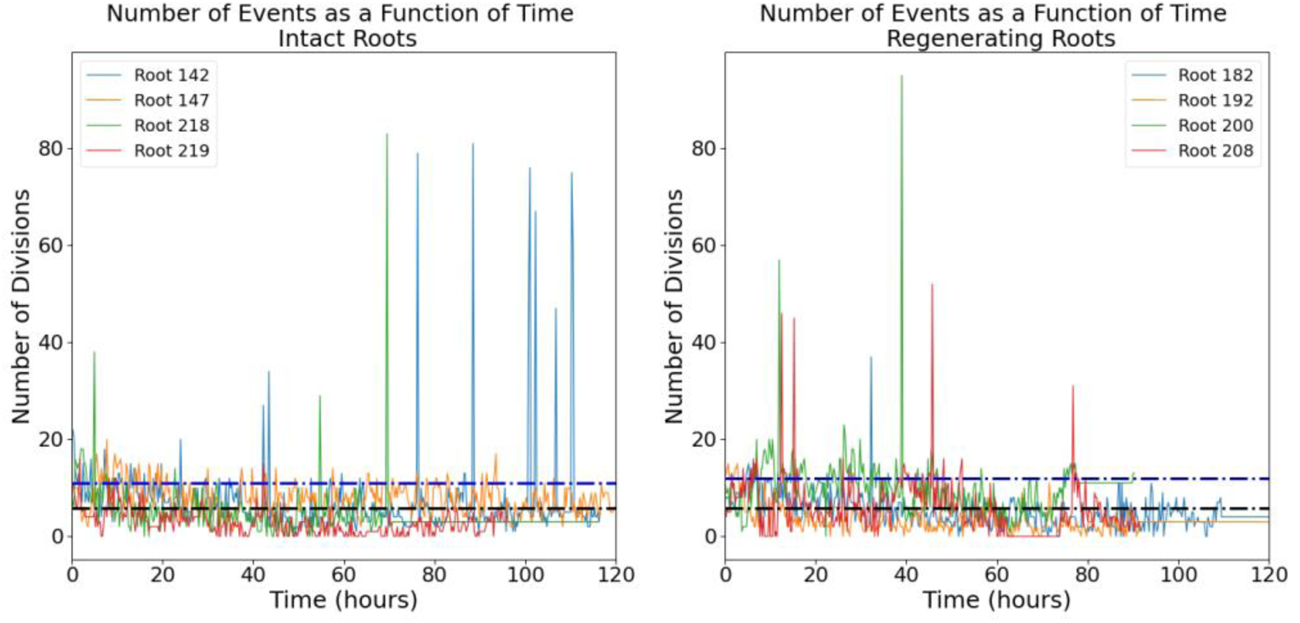
Number of mitotic events detected in intact (left panel) and regenerating (right panel) root tips. Four independent roots are shown for each group. Black dotted line, mean; blue dotted line, mean + standard deviation

### Regenerating and intact roots exhibit different distributions of temporal “bursts” of mitotic events

The intermittent nature of the temporal series in Fig. 1 is interesting and can be further quantified. We define a “burst” as a significant peak in the temporal series. More specifically, a collection of mitotic events occurring in a single time-point and at least one standard deviation higher than the mean of events observed in the entire temporal series. The size of the burst is simply the total number of cell divisions captured at that time point. The two distributions of burst sizes for intact and regenerating roots are significantly different (Fig. 2; K-S test, p<0.01) and indicate that regeneration is on average characterised by larger bursts of cell division activity.

**Figure 2.**
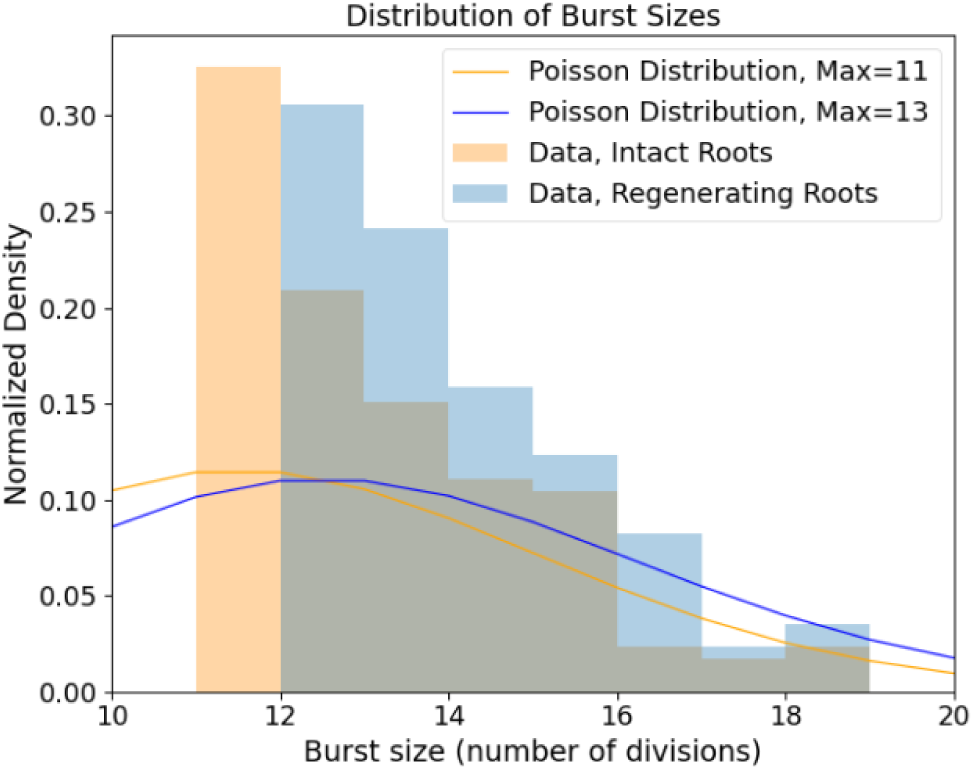
Distribution of burst size (i.e., number of division events in that burst) in intact and regenerating root tips. Experimental data (histograms) and Poisson distributions peaking at 11 (yellow) and 13 (blue) burst sizes.

If the mitotic events were completely uncorrelated from each other, these distributions would be indistinguishable from Poisson distributions. This is not what we observe: The Poisson distribution looks very different from the experimental distribution with the same maximum, both for intact roots and regenerating roots (Fig. 2).

### Regenerating and intact roots exhibit different periodicities of mitotic events

To reveal hidden periodicities in the pattern, we generated a periodogram or a standard spectral analysis of the temporal series of single mitotic events (see Methods). Briefly, periodograms show a distribution of fundamental periodicities in a time series. Our analysis indicates strong fundamental periodicities corresponding to approximately 4, 6 and 24 hours for the intact roots and 11 and 16 hours for the regenerating roots (Fig. 3). Since we enforced a 24-hour light cycle (16-hour light: 8-hour dark) on all the plants during germination, periodicities of 24 hours and its subdivisions (*e*.*g*., 12, 6, 4, etc.) might be expected and trivial. On the other hand, the peak at approximately 16 hours observed in the periodogram of regenerating roots, and not in that of intact roots, suggests a nontrivial periodicity specific to the regeneration process.

**Figure 3.**
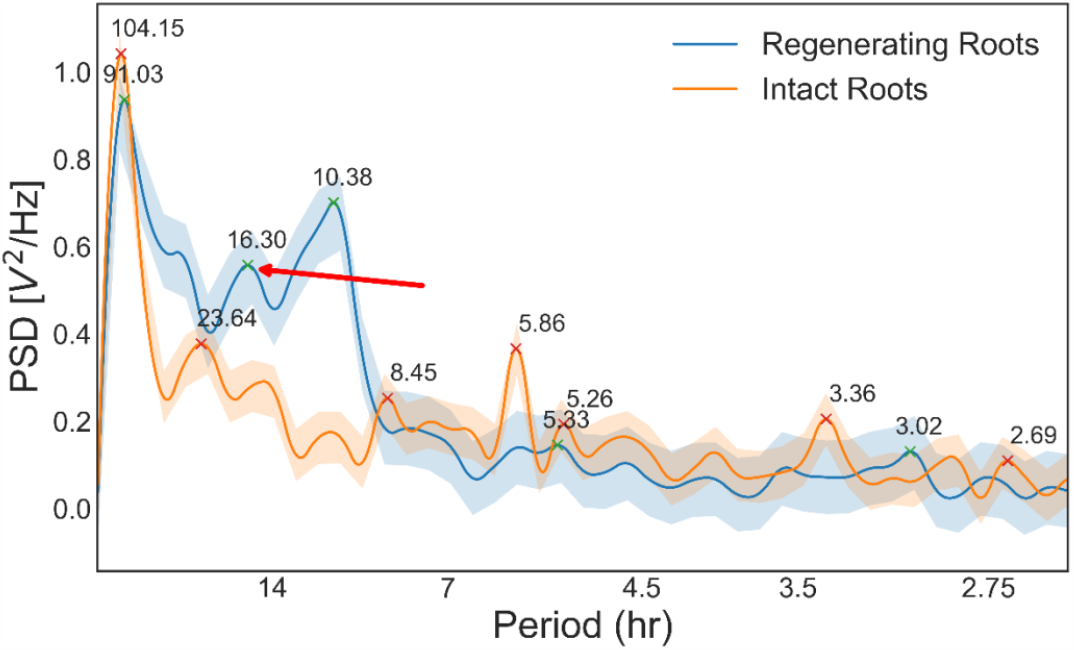
Periodogram of the temporal series shown in Fig. 1 for intact and regenerating root tips. PSD, power spectral density. Red arrow, 16-hour period in regenerating roots, suggesting nontrivial periodicity.

Although the cause of these periodicities remains unclear, the spectral analysis suggests fundamental differences in the cell division dynamics in unperturbed and regenerating tissues.

### A difference in the distribution of mitotic events per frame between regenerating and intact roots emerges only 24 hours after excision

To further characterise the dynamics of mitotic events in both intact and regenerating roots, we compared the distributions of mitotic events in each frame, i.e., the probabilities of detecting a mitotic event at a single time point (Fig. 4). The distributions for intact and regenerating roots are significantly different over the entire duration of our observation (Fig. 4A; K-S test, p<0.001), further supporting the hypothesis that the underlying dynamics of cell divisions are different in intact roots than in regenerating roots.

**Figure 4.**
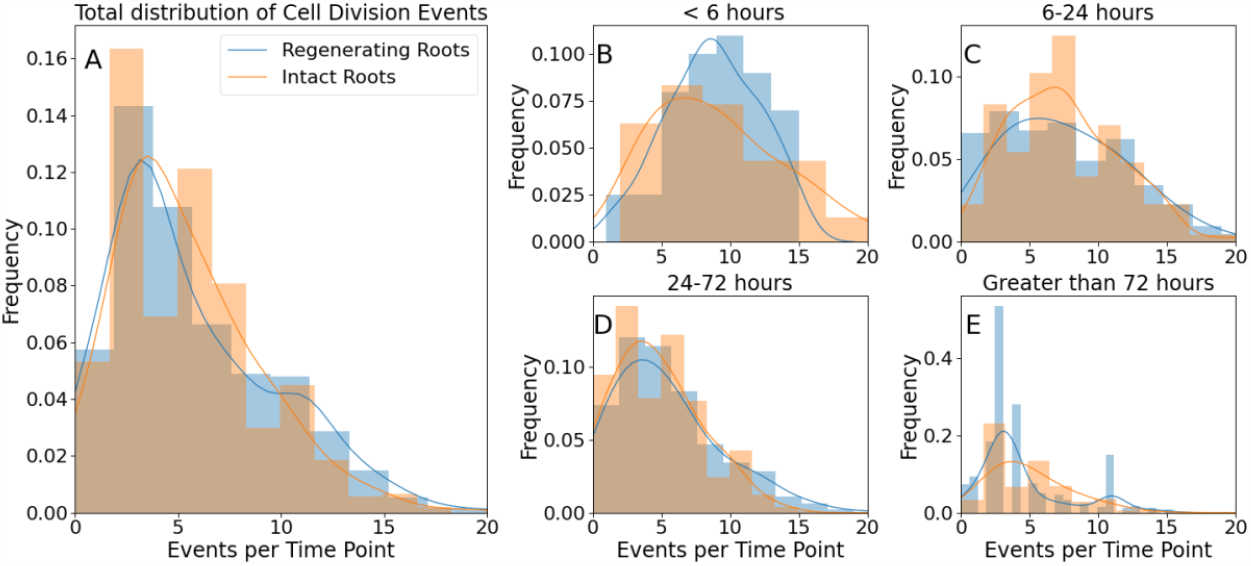
Distributions of mitotic events detected in one frame in intact and regenerating root tips. (A), all events; (B), events detected in the first 6 hours; (C), events detected between 6 and 24 hours; (D), events detected between 24 and 72 hours; (E), events detected after 72 hours. Histograms, experimental data; lines, and kernel density estimation of the experimental data (smooth fitting).

While both distributions peak at approximately 3.5 divisions per frame and are skewed towards higher values, the regenerating root distribution shows a “shoulder” of approximately 11 divisions per frame, which is not as evident in the intact root sample (Fig. 4A). This suggests the existence of two unresolved subpopulations of events in the regenerating roots: one with a maximum of approximately 3.5 divisions per frame, as in the intact roots, and a second one centred at approximately 11 divisions per frame. This second peak is unmatched in the data from the intact roots, suggesting a unique feature of self-organising tissue.

To address whether root regeneration is a single continuous process or, instead, is made of distinct developmental phases, we asked whether the highly active time points with 11 divisions/frame occurred throughout the entire regeneration process or only at specific moments.

We reanalysed the data into temporal bins, 0-6 hours, 6-24 hours, 24-72 hours, and greater than 72 hours after the excision. The distributions of divisions per frame are statistically indistinguishable between intact and regenerating roots during the first 6 hours (Fig. 4B; K-S test, p=0.21) and between 6 and 24 hours (Fig. 4C; K-S test, p=0.83). Crucially, between 24 and 72 hours after excision, the two distributions are marginally significantly different (Fig. 4D; K-S test, p=0.025), with the one for the regenerating roots showing a longer tail between 10 and 20 divisions per frame. Finally, the two distributions remained significantly different 72 hours after excision (Fig. 4E; K-S test, p<0.001). Taken together, these data indicate that the main difference in cell division dynamics between regenerating and intact roots appears only 24 hours after tip excision, with the regenerating roots showing an excess of 10-15 events per time point.

### Mitotic events occur in small spatial clusters that are more abundant in regenerating roots

The lack of a persistent reference point across time frames makes the spatial localisation of the mitotic event relative to biologically significant landmarks in the root intractable. Instead, the spatial information allows the calculation of the relative distance between events. One important question from the developmental point of view is whether these occur uniformly within the tissue or, rather, in clusters.

To define a spatial cluster of events, we first determined the centre of mass of each event using our tracking algorithm (Amarteifio et al., 2021). Around each centre of mass, we modelled a 6 µm X 4 µm X 4 µm cell, with a “diameter” (maximum distance between two points) equal to 8.24 µm. We used the DBScan algorithm (Ester et al., 1996) to identify all events within three cell diameters (ε = 3 x 8.24 µm = 24.72 µm) from each other as part of a single cluster. Finally, we plot the distribution of cluster sizes, or how many clusters of which size we detected at a single time point, for the populations of intact and regenerating roots (Fig. 5). In both distributions, most of the time points contain 1-3 spatial clusters made of 2-4 events each, but at any given time, regenerating roots are more likely to contain a higher number of clusters (up to 4-5) of the same 2-4 cell size (Fig. 5). This can be seen by noting the slightly larger size of the dots at 4-5 clusters in the regenerating roots compared to the same in the intact roots (Fig 5). This subtle distinction suggests a sharp limit in the correlation length among cell division events (i.e., small clusters of cell divisions) but also a propensity of regenerating roots to exhibit a higher number of foci of mitotic activity.

**Figure 5.**
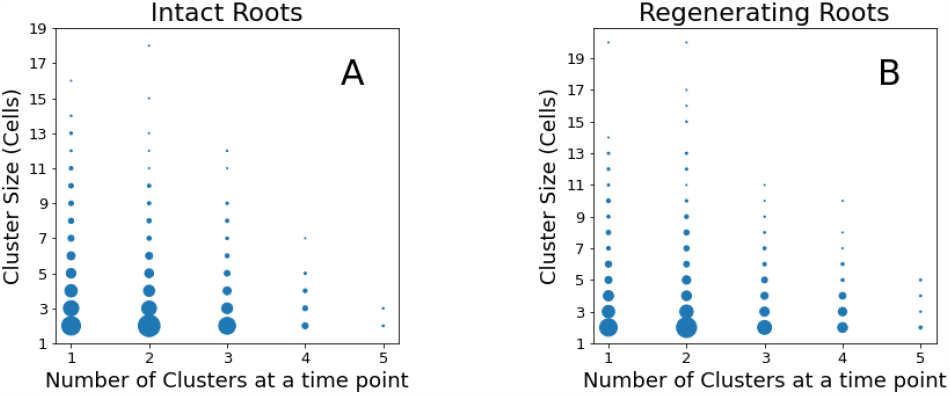
Distribution of cluster numbers and their size at a single time point. The size of each point represents its frequency, or how often that point appears in the data. (A), Intact roots; (B), Regenerating roots.

### The density of mitotic event clusters is constant and analogous between regenerating and intact roots

To quantify the density of cell division clusters in both regenerating and uncut roots, we measured the mean pairwise distance between their centres of mass (Fig. 6). Overall, the two distributions were significantly different (Fig. 6A; K-S test, p<0.001), with a barely significant difference in the first few hours of regeneration (Fig. 6B; K-S test, p=0.001), and then disappeared (Fig. 6C; K-S test, p=0.001) only to become statistically very clear 24 hours after excision (Fig. 6D and 6E; K-S test, p<0.001).

**Figure 6.**
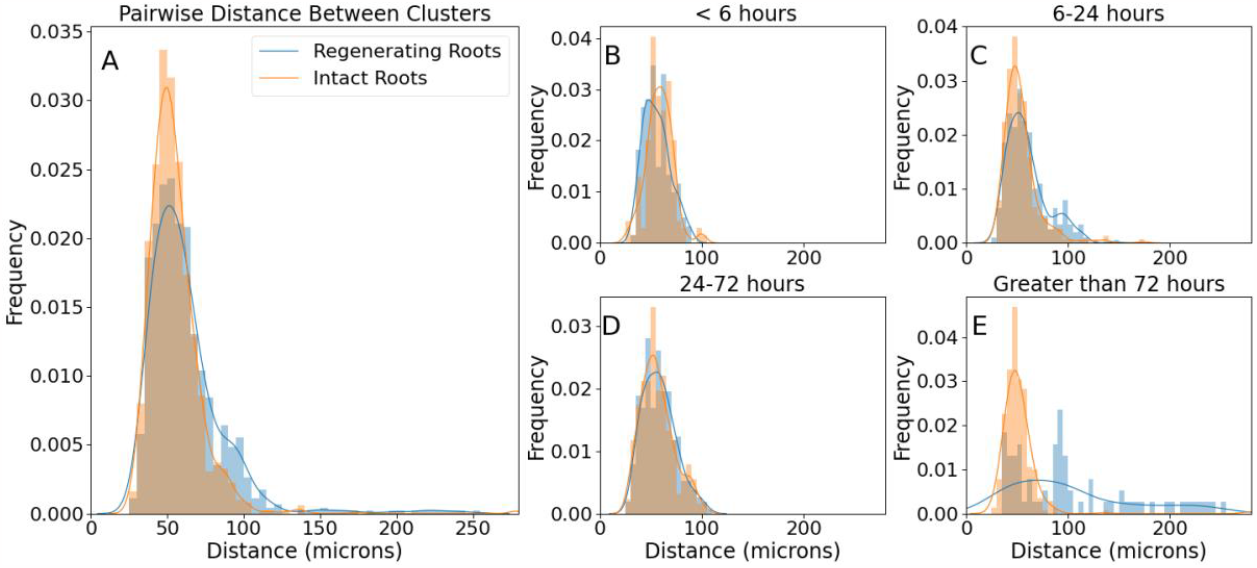
Distributions of pairwise distances between cluster centres of mass. Histograms, experimental data; lines, and kernel density estimation of the experimental data (smooth fitting).

## DISCUSSION

We presented a quantitative characterisation of the temporal and spatial distribution of cell divisions in intact and regenerating *Arabidopsis* root tips. Several biologically relevant observations can be extracted from the data.

First, the intermittent nature of the temporal sequence of mitotic events (Fig. 1) indicates that mitotic events are not randomly distributed in time. In other words, the underlying dynamics of cell divisions in the tissue cannot be explained simply by perfectly uncoupled cells undergoing a noisy cell cycle. A significant body of work describes the complex genetic networks regulating cell-cell interactions during cell division and differentiation in *Arabidopsis* roots, so the fact that cell divisions are not simply independent random events is perhaps not surprising. An intermittent pattern can be described as a sequence of “bursts” or periods of activity above an arbitrary threshold. The distributions of burst size (Fig. 2) look very different than a Poisson distribution, confirming that these are not random, uncorrelated events. This might be expected given the short- and long-range cell-cell signalling, but it is an important quantitative visualisation. Our data also show that regenerating roots tend to produce slightly larger bursts, involving a larger number of cell divisions, compared to intact roots (Fig. 2). This indicates that regeneration entails not only more cell divisions but also that these are compacted in discrete periods (bursts) of higher activity.

Second, cell division activity in both intact and regenerating roots shows a superposition of several periodicities, but regenerating roots are characterised by an underlying period of 16 hours, which is not detected in intact roots (Fig. 3). Although the regenerating and intact groups are composed of random individuals taken from the same isogenic seed population and have been germinated and grown under identical conditions, we note that the seedlings are germinated under a regime of 16 hours in light and 8 hours in darkness. Immediately after root tip excision, the plants were grown and imaged under constant light. Is it possible that a memory of the 16-hour light cycle persists at the cellular level and that it is reflected in the cell division dynamics? If so, our data indicate that this should happen only during tissue regeneration, as no 16-hour periodicity was observed in intact roots. Future experiments carried out with different light/dark regimes might attempt to test this hypothesis.

Third, we found differences between intact and regenerating roots when considering the entire temporal distribution of single mitotic events or the frequency of single time frames containing a given number of cell divisions, despite the described intermittency. More specifically, while in intact roots, cell divisions belong to a single mode centred at approximately 3-5 events at any given time point, during regeneration, a second mode of division emerges, centred at approximately 11 events at any given time point (Fig. 4). This becomes particularly evident 24 hours after root excision, suggesting that after this time point, the regenerating tissue undergoes a transition towards a more complex regime of cell division dynamics. It can be difficult to obtain sufficient temporal and spatial statistics to identify the collective correlations associated with true phase transitions and criticality, so here we limit our reference to a developmental transition.

Fourth, the mitotic events appeared to be clustered in space, suggesting the existence of a short-range inducing signal to trigger cell division in neighbouring cells, coupled with a long-range inhibitory signal to separate clusters. Although it is beyond the scope of this work, we suggest that effective diffusion constants of the inducing and inhibiting signals could be estimated computationally with a model based on reaction-diffusion (Turing, 1952).

Finally, regenerating roots contain a slightly higher number of clusters per frame (Fig. 5), which are also more densely distributed (i.e., with smaller inter-cluster distance) when compared to intact roots (Fig. 6). This suggests that a similar tissue volume is going through cell proliferation in regenerating and intact roots but that subregions of high mitotic activity (clusters) appear more often in the regenerating tissue.

Overall, the presented data paint an original quantitative picture where the cell division dynamics in regenerating roots evolve faster than those in intact roots, possibly revealing a developmental transition approximately 24 hours after physical perturbation. Although this is only a first step towards a full quantitative characterization of tissue regeneration, we believe that the focus on cell divisions is important to capture the complex dynamics driving tissue self-organization.

## METHODS

### Plant material

Mitotic events were visualised using an existing Arabidopsis transgenic line expressing the cyclin-GFP fusion CYCB1;1::GFP (Reddy et al., 2004). Arabidopsis seeds were sterilised, stratified and stored at 4°C before sowing on sterile room temperature rectangular plates prepared in sterile conditions with solid media consisting of 0.175% w/v Murashige and Skoog Basal Medium (MS) (Sigma-Aldrich, UK), 0.5% w/v sucrose (Sigma-Aldrich, UK), 0.05% w/v MES hydrate (Sigma-Aldrich, UK), and 0.8% w/v agar, adjusted to pH 5.7 (KOH), which was sterilised by autoclaving. The plates were placed in vertical racks in a plant growth chamber with 120 μmol/m2/s light intensity on a 16:8 hour light:dark cycle and constant 23°C.

### Microdissection

Regenerating roots were manually excised using a 100 Sterican 27G needle tip (B Braun) under a Nikon SMZ1000 dissecting microscope, 180x magnification, following published procedures (Kral et al., 2016; Sena et al., 2009). The excisions were mainly performed at ∼100 µm, with one root being excised at ∼50 µm.

### Mounting

Five days post-germination, plants were moved and mounted in an imaging cuvette as previously described (Baesso et al., 2018). Briefly, roots were taken and placed on solid media plates with 5% w/v agar (all other reagents were the same as the germination plates).

Excised and control roots were then both mounted into the corner of an imaging cuvette (manufacturer) by flowing liquid media (0.04375% w/v MS, 0.5% w/v sucrose, 0.05% w/v MES) and using capillary action to pull the root down the length of the cuvette with the hypocotyl and cotyledons above the top of the cuvette. The root was held in place with a sterile, heat-shrink plastic-coated pin, which was in turn held in place by 2 mm glass beads for 1/3 of the volume of the cuvette, followed by 1 mm glass beads until 10 mm from the top of the cuvette. Liquid media was perfused into the chamber at ∼1 mL/min through a custom cuvette top with a recessed corner for the cotyledons. A second cuvette with a glass coverslip top and two ∼0.5 cm^2 gas-exchange windows covered with gas-permeable sterile tape was placed over the top of the perfusion chamber, allowing for gas exchange and broad-spectrum incident light on the cotyledons. Media temperature was monitored using an infrared thermometer mounted on tubing before the perfusion chamber, and a multistage heating element was used to keep the media temperature at 23°C. All tubing, media, glass beads, pins, perfusion tops and imaging cuvettes were autoclaved before use. Plastic imaging chamber tops were submerged in a bleach solution for 30 s before multiple rinses with autoclaved sterile water.

### Microscopy

Imaging was performed on a previously described home-built light-sheet microscope (Baesso et al., 2018). The root was imaged through 60 planes every 15 minutes for up to 7 days. At each plane, 6 images were taken, and the plane of maximum focus was kept. Focusing was enhanced by automatically detecting the edge of the root at the beginning of every image set, moving the focus 30 µm into the root from that plane, automatically detecting the plane of maximum focus, and moving back 30 µm from that point, such that the starting point for the autofocusing step would be at maximum focus in the centre of the root. The root tip is automatically tracked using custom MATLAB code (Baesso et al., 2018), which in turn will move the cuvette stage in x, y, and z to keep the root in focus and centred in the field of view throughout the experiment. Cell division events were segmented and tracked across time frames using a previously described Python code (Amarteifio et al., 2021).

### Statistical analysis

When comparing two samples of measurements, the nonparametric two-sample Kolmogorov-Smirnov (KS) test was used. Unless stated otherwise, all comparisons were performed assuming independence (unpaired test). All statistical tests were performed in Python using the *scipy* statistics package, and the analysis described in Jupyter Notebooks stored at https://github.com/GiovanniSena/Fallesen_2024.

## AUTHORSHIP CONTRIBUTIONS

T.F. performed the experiments, analysed the data (including statistics) and wrote the article; S.A., G.P. and H.J.J. all contributed to the analysis of the data; G.S. conceived and designed the project, supervised the experiments and contributed to the writing of the manuscript. All authors revised and approved the final manuscript.

## DATA AVAILABILITY STATEMENT

The images generated in this study are available from the corresponding author upon request.

## REFERENCES

Amarteifio, S., Fallesen, T., Pruessner, G., & Sena, G. (2021). A random-sampling approach to track cell divisions in time-lapse fluorescence microscopy. Plant Methods, 17(1), 25. 10.1186/s13007-021-00723-8

Baesso, P., Randall, R. S., & Sena, G. (2018). Light Sheet Fluorescence Microscopy Optimized for Long-Term Imaging of Arabidopsis Root Development. Methods in Molecular Biology (Clifton, N.J.), 1761(7), 145–163. 10.1007/978-1-4939-7747-5_11

Balaguer, M. A. de L., Ramos-Pezzotti, M., Rahhal, M. B., Melvin, C. E., Johannes, E., Horn, T. J., & Sozzani, R. (2016). Multi-sample Arabidopsis Growth and Imaging Chamber (MAGIC) for long term imaging in the ZEISS Lightsheet Z.1. Developmental Biology, 419(1), 19–25. 10.1016/j.ydbio.2016.05.029

Beemster, G. T. S., & Baskin, T. I. (1998). Analysis of Cell Division and Elongation Underlying the Developmental Acceleration of Root Growth in Arabidopsis thaliana. Plant Physiology, 116(4), 1515–1526. 10.1104/pp.116.4.1515

Berthet, B., & Maizel, A. (2016). Light sheet microscopy and live imaging of plants. Journal of Microscopy, 263(2), 158–164. 10.1111/jmi.12393

Buckner, E., Madison, I., Chou, H., Matthiadis, A., Melvin, C. E., Sozzani, R., Williams, C., & Long, T. A. (2019). Automated Imaging, Tracking, and Analytics Pipeline for Differentiating Environmental Effects on Root Meristematic Cell Division. Frontiers in Plant Science, 10, 1487. 10.3389/fpls.2019.01487

Campilho, A., Garcia, B., Toorn, H. V. D., Wijk, H. V., Campilho, A., & Scheres, B. (2006). Time-lapse analysis of stem-cell divisions in the Arabidopsis thaliana root meristem. The Plant Journal : For Cell and Molecular Biology, 48(4), 619–627. 10.1111/j.1365-313x.2006.02892.x

Clark, N. M., Broeck, L. V. den Guichard, M., Stager, A., Tanner, H. G., Blilou, I., Grossmann, G., Iyer-Pascuzzi, A. S., Maizel, A., Sparks, E. E., & Sozzani, R. (2020). Novel Imaging Modalities Shedding Light on Plant Biology: Start Small and Grow Big. Annual Review of Plant Biology, 71(1), 1–28. 10.1146/annurev-arplant-050718-100038

Dinsmore, C. E. (1991). A History of regeneration research : milestones in the evolution of a science. Cambridge University Press.

Elchaninov, A., Sukhikh, G., & Fatkhudinov, T. (2021). Evolution of Regeneration in Animals: A Tangled Story. Frontiers in Ecology and Evolution, 9, 621686. 10.3389/fevo.2021.621686

Ester, M., Kriegel, H.-P., Sander, J., & Xu, X. (1996). A density-based algorithm for discovering clusters in large spatial databases with noise. Proceedings of the Second International Conference on Knowledge Discovery and Data Mining, 226–231.

Ikeuchi, M., Ogawa, Y., Iwase, A., & Sugimoto, K. (2016). Plant regeneration: cellular origins and molecular mechanisms. Development, 143(9), 1442–1451. 10.1242/dev.134668

Kral, N., Ougolnikova, A. H., & Sena, G. (2016). Externally imposed electric field enhances plant root tip regeneration. Regeneration, 3(3), 156–167. 10.1002/reg2.59

Lavrekha, V. V., Pasternak, T., Ivanov, V. B., Palme, K., & Mironova, V. V. (2017). 3D analysis of mitosis distribution highlights the longitudinal zonation and diarch symmetry in proliferation activity of the Arabidopsis thaliana root meristem. The Plant Journal : For Cell and Molecular Biology, 92(5), 834–845. 10.1111/tpj.13720

Liu, X., Zhu, K., & Xiao, J. (2023). Recent advances in understanding of the epigenetic regulation of plant regeneration. aBIOTECH, 4(1), 31–46. 10.1007/s42994-022-00093-2

Maizel, A., Wangenheim, D. von, Federici, F., Haseloff, J., & Stelzer, E. H. K. (2011). High-resolution live imaging of plant growth in near physiological bright conditions using light sheet fluorescence microscopy. The Plant Journal : For Cell and Molecular Biology, 68(2), 377–385. 10.1111/j.1365-313x.2011.04692.x

Morgan, T. H. (1901). Regeneration. The Macmillan Company; Macmillan & Co. ltd.

Morinaka, H., Sakamoto, Y., Iwase, A., & Sugimoto, K. (2023). How do plants reprogramme the fate of differentiated cells? Current Opinion in Plant Biology, 74, 102377. 10.1016/j.pbi.2023.102377

Rahni, R., & Birnbaum, K. D. (2019). Week-long imaging of cell divisions in the Arabidopsis root meristem. Plant Methods, 15(1). 10.1186/s13007-019-0417-9

Reddy, G. V., Heisler, M. G., Ehrhardt, D., & Meyerowitz, E. M. (2004). Real-time lineage analysis reveals oriented cell divisions associated with morphogenesis at the shoot apex of Arabidopsis thaliana. Development, 131(17), 4225–4237. 10.1242/dev.01261

Sena, G., Frentz, Z., Birnbaum, K. D., & Leibler, S. (2011). Quantitation of cellular dynamics in growing Arabidopsis roots with light sheet microscopy. PLoS One, 6(6), e21303. 10.1371/journal.pone.0021303

Sena, G., Wang, X., Liu, H.-Y., Hofhuis, H., & Birnbaum, K. D. (2009). Organ regeneration does not require a functional stem cell niche in plants. Nature, 457(7233), 1150–1153. 10.1038/nature07597

Sugimoto, K., Temman, H., Kadokura, S., & Matsunaga, S. (2019). To regenerate or not to regenerate: factors that drive plant regeneration. Current Opinion in Plant Biology, 47, 138–150. 10.1016/j.pbi.2018.12.002

Turing, A. M. (1952). The Chemical Basis of Morphogenesis. Philos T Roy Soc B, 237(641), 37–72. http://www.google.com/search?client=safari&rls=en-us&q=The+Chemical+Basis+of+Morphogenesis&ie=UTF-8&oe=UTF-8

Wangenheim, D. von, Fangerau, J., Schmitz, A., Smith, R. S., Leitte, H., Stelzer, E. H. K., & Maizel, A. (2016). Rules and Self-Organizing Properties of Post-embryonic Plant Organ Cell Division Patterns. Current Biology, 26(4), 439–449. 10.1016/j.cub.2015.12.047

